# Using RNASeq to investigate the involvement of the *Ophiocordyceps* clock in ant host infection and behavioral manipulation

**DOI:** 10.1101/2023.01.20.524843

**Authors:** Biplabendu Das, Ian Will, Roos Brouns, Andreas Brachmann, Charissa de Bekker

## Abstract

**Introduction:** Parasites can modify host behavior to ensure their own growth and transmission. Multiple species of the fungi *Ophiocordyceps* infect ants, but in a species-specific manner; one fungal species co-evolved to successfully modify the behavior of one ant species. However, several characteristics of the behavioral modification seem to be similar across different *Ophiocordyceps*-ant systems, including a preference for the time of the day for manipulating host behavior. In this study, we explored the various mechanisms via which the circadian clock of *Ophiocordyceps* might be playing a role in modifying host behavior. We studied *O. camponoti-floridani* that modifies the behavior of its ant host *Camponotus floridanus*. To separate the role of the clock in behavior manipulation, from its role in growth and survival, we compared the daily gene expression profile of *O. camponoti-floridani* to a generalist, non-manipulating fungal parasite, *Beauveria bassiana*, which also successfully infects the same ant host.

**Results:** Majority of the 24h rhythmic *O. camponoti-floridani* genes show peak expression before or at the transitions between light and dark. Rhythmic genes in *O. camponoti-floridani*, for which *B. bassiana* lacks an ortholog, were overrepresented for enterotoxin genes. Around half of all genes that show 24h rhythms in either *O. camponoti-floridani* or *B. bassiana* showed a consistent difference in their temporal pattern of daily expression. At the halfway mark in *O. camponoti-floridani* infections, when diseased ants show a loss of 24h rhythms in daily foraging, several fungal clock genes, including *Frequency*, showed differential expression. Network analyses revealed a single gene cluster, containing *White Collar 1* and *2*, that showed overrepresentation for genes oscillating every 24h in liquid culture as well as genes differentially expressed while growing inside the ant head.

**Conclusion:** This study identifies several sets of putatively clock-controlled genes and biological processes in *O. camponoti-floridani* that likely plays a role in modifying the behavior of its ant host. Differential expression of *O. camponoti-floridani* clock genes or 24h-rhythmic genes during infection is suggestive of either a loss of daily rhythm or a change in the amplitude of rhythmic gene expression. Both possibilities would suggest that a disease-associated change occurs to the functioning of the *O. camponoti-floridani* clock, and its output, while the fungi grows inside the ant head.

## INTRODUCTION

Several parasites have evolved the ability to adaptively change their host’s behavior to facilitate their survival and transmission. Examples of manipulating parasites and their hosts are found across several taxa [1–3]. Among parasitic manipulators, the *Ophiocordyceps* fungi that infect ants are emerging as a powerful model system. *Ophiocordyceps*-ant systems are highly species-specific; one species of *Ophiocordyceps* can successfully manipulate the behavior of only one species of ant. This is consistent with the fact that these parasite-host pairs have co-evolved for millions of years [4]. Although these parasite-host systems show species-specificity, the disease phenotype is similar across all *Ophiocordyceps*-ant systems. In the final stages of the disease, *Ophiocordyceps* infected ants tend to show stereotypical behavioral changes that are adaptive to the parasite: social isolation from the colony, phototactic summiting, and eventual attachment at vantage points that promote fungal development and transmission.

The manipulated biting seems highly synchronized to a specific time of day. Field observations of an *Ophiocordyceps*-carpenter ant system in Thailand showed that the biting happens primarily around solar noon [5]. A similar time-of-day synchronized biting was observed in controlled lab studies with two completely different *Ophiocordyceps*-ant pairs [6, 7]. The conserved nature of timely host manipulation across *Ophiocordyceps*-ant pairs strongly suggests an underlying role of biological clocks [8].

Our findings from a related study demonstrated that at least the host’s clock is involved [9]. For example, at the halfway mark in the disease, *Camponotus floridanus* ants infected with *O. camponoti-floridani* showed synchronized changes in the daily expression of several clock-controlled genes previously found to show caste-associated differential rhythmicity [9]. Such alterations to the host’s rhythmic processes have also been found in other manipulating parasite-host systems. For example, baculoviruses are known to manipulate infected caterpillars to show a phototactic summiting behavior before death. Phototactic behavior in infected caterpillars seems to be correlated with altered host phototransduction and circadian entrainment pathways [10, 11].

What remains less clear is the precise role of the parasite’s clock in inducing timely changes to host rhythms. A previous study by de Bekker and colleagues has shown that *Ophiocordyceps*, like most fungi, has homologs of known core clock components that make up the transcription-translation feedback loop [12]. Using time-course RNASeq, the authors also showed that the daily expression of several secreted enzymes, proteases, toxins, and small bioactive compounds show robust 24h-rhythms. However, it is yet unknown to what degree the disease outcome, especially changes to host’s clock-controlled processes, is driven by the parasite’s clock. If the clock of *Ophiocordyceps* is important for inducing timely manipulation of its host ant’s behavior, it would be worth investigating how the clock of a manipulating fungus functionally differs from that of other fungal entomopathogens that do not manipulate host behavior. The latter question is important to address because, although studies of plant pathogens have demonstrated the role of fungal clocks in regulating virulence, we do not know much about the role clocks play in the success of fungal entomopathogens in general, let alone for behavior modifying fungi [13, 14]. Therefore, to understand the mechanisms via which the *Ophiocordyceps* clock might enable manipulation of ant behavior, we need to disentangle the role of fungal clocks in modifying host behavior from its role in the parasite’s growth and survival. This study aimed to characterize the clock and clock-controlled processes of the behavior-manipulating fungi *Ophiocordyceps camponoti-floridani* compared to a generalist, non-manipulating entomopathogen, *Beauveria bassiana*, to shed light on the role of fungal clocks in inducing timely changes to host’s clock-controlled processes, including foraging behavior.

Compared to *O. camponoti-floridani*, *B. bassiana* induces a vastly different disease progression inside *C. floridanus* ants. A clear difference can be found in the survival times of the infected host; in controlled lab conditions, ants infected with *O. camponoti-floridani* can live up to three to four weeks, whereas *B. bassiana*-infected ants die within a week [15, 16]. Since *B. bassiana* functions as a generalist parasite that kills and consumes its host within a matter of days upon infection, without the need to induce pathogen-adaptive behavior in its host, its lifestyle can be classified as necrotrophic. On the contrary, *O. camponoti-floridani*, which co-evolved with and highly specialized to infect one host, *C. floridanus*, spends more time in a seemingly symbiotic relationship with its host before killing it. Such a parasitic lifestyle can be classified as hemibiotrophic (reviewed in [17]). Here we discuss the similarities and differences in the functioning of the biological clock of *B. bassiana* as compared to *O. camponoti-floridani*. The comparison between *B. bassiana* and *O. camponoti-floridani* fits multiple, non-mutually exclusive contexts: (1) a necrotroph versus a hemibiotroph, (2) a generalist versus a specialist, and finally, (3) a non-manipulating versus behavior manipulating parasite.

Using time-course RNASeq and comparative network analyses, we compared the daily transcriptome of *Ophiocordyceps camponoti-floridani* and *Beauveria bassiana* (synonymous with *Cordyceps bassiana*), both of which are known to successfully infect the Florida carpenter ant *Camponotus floridanus* [18]. More specifically, we characterized the similarities and differences in how the clocks of these two fungal parasites temporally segregate their biological processes as they grow in their blastospore stage in controlled liquid media outside the ant host. We also discuss how the manipulating parasite *O. camponoti-floridani*’s clock-controlled gene expression changes while it grows inside the ant head at the halfway mark in disease progression. To do so, we used network analyses to identify potential regulatory mechanisms via which the clock of *O. camponoti-floridani* might interact with the host’s clock or regulate aspects of its own timekeeping as it grows inside the host’s head. The corresponding changes to the host *C. floridanus*’s daily transcriptome during infection induced by both fungal entomopathogens, at the halfway mark in disease progression, has been documented in a separate research article [9].

## MATERIALS AND METHODS

### Fungal culturing and circadian entrainment

To culture *O. camponoti-floridani* and *B. bassiana*, we used the same protocol and fungal strains (Arb2 and BB0062, respectively) described in [6, 16]. To reiterate the process briefly, we grew both *O. camponoti-floridani* and *B. bassiana* in their blastospore stage in Grace’s Insect Medium (Gibco, Thermo Fisher Scientific) supplemented with 2.5% FBS (Gibco) using established protocols (de Bekker et al., 2017; Will et al., 2020; Ying & Feng, 2006). As for entraining fungal samples for time-course RNASeq, we followed the protocol described in de Bekker et al. (2017). We kept the fungal cultures under strict 12h:12h light-dark cycles while setting the temperature and relative humidity constant. To ensure that the fungi were entrained to the new light-dark regime readily, we kept fungal cultures under constant light conditions for 48 hours (constant temperature and relative humidity) before moving them to strict 12h:12h light-dark cycles. We allowed the fungal cultures to entrain to light-dark cycles for five days before harvesting them for RNASeq. To perform time-course RNASeq, however, we needed to harvest fungal blastospores at multiple time points throughout the day. We split the original (source) culture undergoing entrainment for each fungal species into 18 separate fungal cultures (12 for sampling and 6 as backup) on day four of light-dark entrainment. We reseeded 250 μL of source fungal culture (OD at 660nm = 1.089) into 15 mL of fresh media. Providing blastospores with fresh media also ensured that the fungal cells didn’t experience starvation during sampling. Following the split, all fungal cultures demarcated for sampling were left undisturbed under the light-dark entrainment conditions until sampling day.

### Harvesting fungal samples over a day under 12:12 LD

We harvested both fungal samples simultaneously on day six of the light-dark entrainment period. Starting at ZT2 (two hours after lights were turned on), we harvested fungal blastospores every 2h over a 24h period. At a given sampling time point, for each species, we pelleted the fungal cells by spinning down 2 mL of the fungal culture for 3 min at 10,000 rpm. The supernatant was discarded, and the process was repeated twice for *O. camponoti-floridani* and thrice for *B. bassiana* to obtain a comparable size of fungal pellets (condensed blastospores). We then added two ball bearings to each pellet, flash-froze the samples in liquid nitrogen, and kept them at −80 °C until further processing for RNASeq. We obtained twenty-four fungal samples from the two species, harvested across twelve time points throughout the 24h day.

### RNA extraction, library preparation, and RNASeq

We extracted total RNA from the frozen fungal samples and prepared the cDNA libraries for RNASeq using the same protocol described in [6]. All twenty-four cDNA libraries were sequenced as 50 bp single-end reads on an Illumina HiSeq1500 at the Laboratory for Functional Genome Analysis (Ludwig-Maximilians-Universitat Gene Center, Munich).

### Pre-processing sequencing data

We first removed sequencing adapters, and low-quality reads from our RNASeq data using BBDuk (parameters: qtrim = rl, trimq = 10, hdist = 1, k = 23) [19]. Post-trimming, we used HISAT2 [20] to map transcripts to the relevant genomes (*O. camponoti-floridani* (NCBI ID: 91520; Will et al. 2020) and *B. bassiana* (ARSEF 2860; Xiao et al. 2012)). Finally, we obtained normalized gene expression from the mapped reads as Fragments Per Kilobase of transcript per Million (FPKM) using Cuffdiff [21].

### Rhythmicity and Overrepresentation analyses

To perform functional enrichment analyses, we used an updated version of the custom enrichment function that performs hypergeometric tests (previously used in [6] and [22]). The function “check_enrichment” is available via the timecourseRnaseq package on GitHub (https://github.com/biplabendu/timecourseRnaseq) [23]. For a given set of genes, we identified overrepresented Gene Ontology (GO) terms or PFAM domains using the check_enrichment function, and significance was inferred at 5% FDR. If not mentioned otherwise, for functional enrichment tests, we used all genes that were found to be “expressed” (≥ 1 FPKM expression for at least one sample) in the ant heads for each treatment as the background gene set. We only tested GO/PFAM terms annotated for at least five protein-coding genes. To cluster a set of genes according to their daily expression, we used an agglomerative hierarchical clustering framework (method = complete linkage) using the ‘hclust’ function in the stats package for R. To determine differentially expressed genes, we used the linear modeling framework proposed in LimoRhyde [24], but without interaction between treatment and time. A gene was considered significantly differentially expressed if treatment was found to be a significant predictor (at 5% FDR), and the gene showed at least a two-fold change in mean diel expression between the two treatment groups (abs(log_2_-fold-change) ≥ 1).

All data analyses, including statistical tests and graphical visualizations, were performed in RStudio [25] using the R programming language v3.5.1 [26]. Heatmaps were generated using the pheatmap [27] and viridis [28] packages. Upset diagrams were used to visualize intersecting gene sets using the UpsetR package [29]. We used Fisher’s exact test to identify if two genesets showed significant overlap (significance was inferred at 5% FDR). We visualized the results of multiple pairwise Fisher’s exact tests using the GeneOverlap package [30]. All other figures were generated using the ggplot2 package [31].

### Differentially expressed O. camponoti-floridani genes during infection

Although blastospores represent the fungal life stage in which it exists inside infected ant hosts, we expect the fungal gene expression profiles to differ when it grows inside the host’s body. This is because once inside the ant host, the fungal pathogen is subjected to different entrainment cues mediated by the host’s physiology and a newfound competition against the host’s microbiota and immune repertoire. In a separate study [9], we performed RNA-Sequencing on *O. camponoti-floridani*-infected ant heads collected halfway through its disease progression using the same sampling resolution and light-dark entrainment conditions used in this study to harvest fungal blastospores. At the midpoint in disease progression, ants infected with *O. camponoti-floridani* appear to lose their daily activity rhythms [9, 16]. At every 2h, over a 24h period, we pooled extracted RNA from three infected ant heads to prepare the cDNA libraries for sequencing (see Methods in [9] for more details). Performing RNASeq on the entire ant head, instead of the brain, allowed us to obtain a mixed transcriptome containing both ant and *O. camponoti-floridani* mRNA and, eventually, normalized gene expression data for both host and parasite.

Although, on average, we obtained 8.1 (± 1.8) million reads per sample (timepoint) that mapped uniquely to the *O. camponoti-floridani* genome, the pooled sample we collected at ZT6 displayed a relatively low read depth (1.1 million reads). However, sufficient sequencing reads for each sampling timepoint were necessary to confidently characterize the changes to the fungal daily gene expression profiles during disease. Consequently, we could not confidently run the rhythmicity analyses for *O. camponoti-floridani* daily transcriptomes during disease since the sharp peaks or troughs we observe in a gene’s expression at ZT6 could easily be due to the low sequencing depth of the sample. Therefore, we decided against interpreting the changes to rhythmic properties of fungal gene expression in ant heads at the halfway mark in disease.

However, we were able to quantify disease-associated increases or decreases in the expression (differential expression) of *O. camponoti-floridani* genes. We used a linear modeling framework proposed in LimoRhyde [24] to classify genes as differentially expressed, but without any interaction term. A gene was considered differentially expressed if the treatment was found to be a significant predictor (at 5% FDR), and there was at least two-fold difference in expression levels of the *O. camponoti-floridani* gene at the halfway mark in disease as compared to growth outside the host.

### Network analyses

To build the annotated gene co-expression network (GCN), we used the exact protocol detailed in de Bekker and Das (2022) [8]. The clustering of genes into co-expressed modules was performed using functions from the WGCNA package [32–34], the gene-gene and module-module correlations were calculated using Kendall’s tau-b correlation [35], and the global connectivity patterns of the GCN was visualized using the igraph package [36]. The significance of pairwise overlaps was calculated using Fisher’s exact tests and visualized using the GeneOverlap package [30].

## RESULTS AND DISCUSSION

### General patterns of daily gene expression in both fungal entomopathogens

To identify the putative genes and processes under clock control in *O. camponoti-floridani* and *B. bassiana* and to characterize the differences in their daily timekeeping, we performed RNA-Seq on 12h:12h light-dark entrained fungal cultures that were harvested every 2h, over a 24h period. For each time point, we collected the two fungal species simultaneously in their yeast-like blastospore stage, which is the primary life stage of the fungi inside their infected ant host (Wang and Wang 2017).

We classified fungal genes as “expressed” if their mRNA levels were greater than 1 FPKM for at least one of the 12 sampling time points during the 24h period. Of the 7455 protein-coding genes annotated in the *O. camponoti-floridani* genome, 94% (6998 genes) were expressed, whereas only 2.5% (190 genes) showed no expression (FPKM = 0 at all time points). As for *B. bassiana*, 9006 (87%) of the 10364 protein-coding genes were expressed at some point during the 24h day, and 756 (7.3%) were not expressed. The non-expressed genes for *O. camponoti-floridani*, but not *B. bassiana*, showed an overrepresentation of heat-labile enterotoxin genes (five out of the 34 annotated in the genome) putatively involved in pathogenesis and interspecies interaction between organisms (Fig. 1), of which some have been hypothesized to be involved in behavioral manipulation (de Bekker et al., 2015; de Bekker et al., 2017; Will et al., 2020). Among these five *O. camponoti-floridani* genes, two copies of heat-labile enterotoxin A subunit did not show any expression (FPKM = 0) even inside ant hosts at halfway through disease progression (also referred to as “infection” throughout this manuscript). In contrast, two putative enterotoxins and a FAD-binding domain-containing protein showed only a low expression (0 < FPKM < 1) during infection.

**Figure 1:**
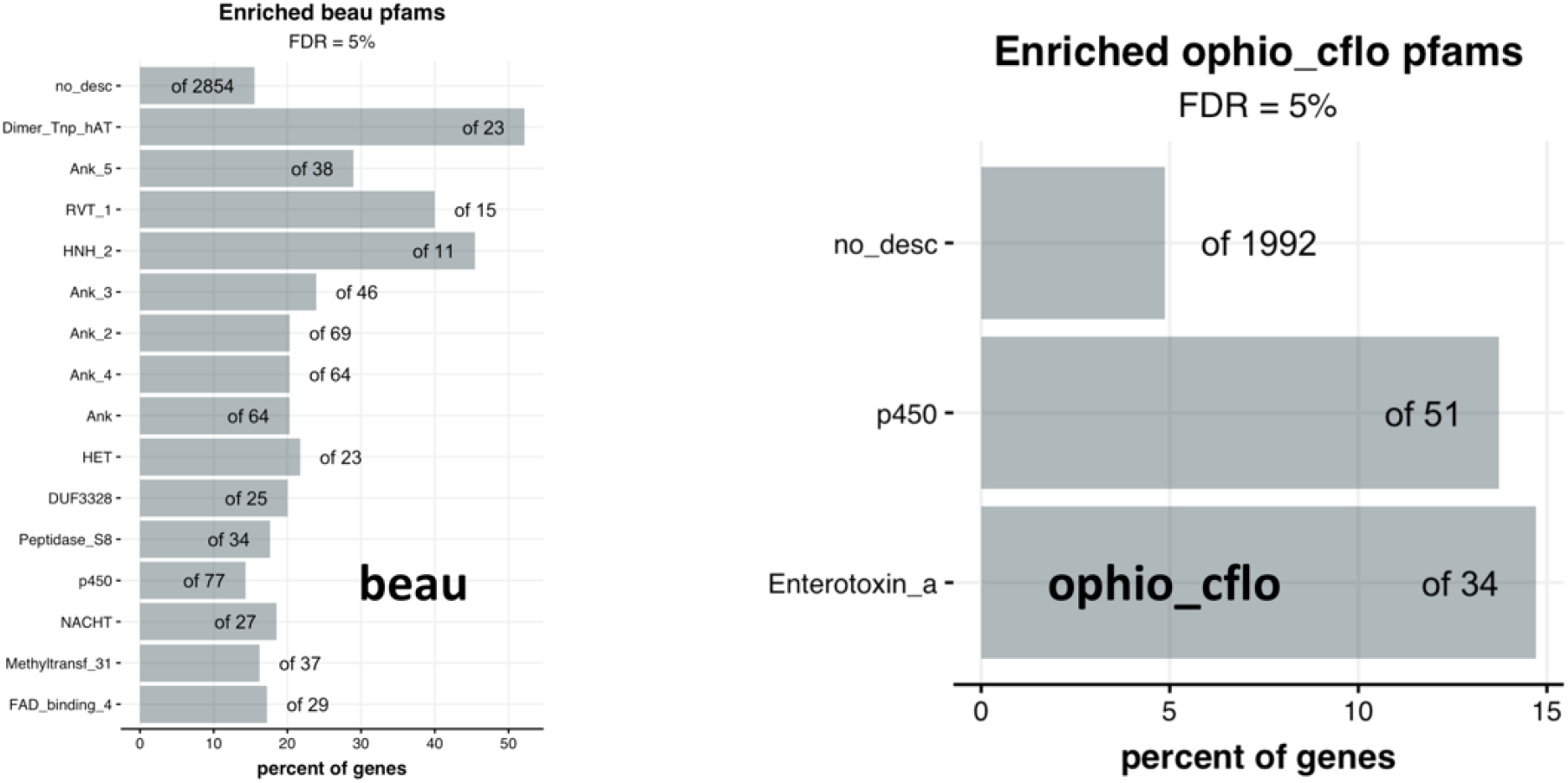
Functions of fungal genes not expressed in their blastospore stage while growing in controlled conditions outside the host body. The two panels show the PFAM domains overrepresented in the set of genes that show no expression (FPKM=0) in (left) *B. bassiana* and (right) *O. camponoti-floridani* during their blastospore growth stage in controlled conditions. The bars represent the percent of genes annotated with a given pfam domain found in the test gene set (in this case, non-expressed genes) compared to all such genes found in the background gene set (in this case, all genes in the ant genome). Unless mentioned otherwise, all subsequent enrichment plots used in this manuscript have the same meaning.

### Daily rhythms in gene expression – O. camponoti-floridani

Of the 6889 *O. camponoti-floridani* genes expressed in their blastospore stage, 33% (2285 genes) showed significant 24h-rhythms in daily expression. We used hierarchical clustering of these rhythmic genes to tease apart clusters of genes that show the same (or similar) phase of daily expression. We divided the rhythmic genes into four clusters upon hierarchical clustering to potentially identify day, night, dusk, and dawn peaking genes (Fig. 2A-B). We identified 354 genes (Fig. 2A-B, cluster-3) that showed clear day-peaking activity, with more than 50% (172 genes) of genes oscillating with a phase at ZT2, two hours after lights were turned on. It is unclear if the peak activity at ZT2 for these processes is driven by the endogenous clock (anticipatory) or if it is a response to light (reactive) since the transition from dark to light happens two hours earlier, at ZT24 (ZT24 is the same as ZT0). Next, we identified another set of 633 putative day-active genes (cluster-4) that showed a more distributed phase of daily expression as compared to cluster-3 genes (Fig. 2A). The majority of cluster-4 genes (45% of 633 genes) showed a peak expression between ZT10 and ZT12. The latter – ZT12 – is the time at which the fungal culture experienced light to dark transition. Before and during harvesting samples for RNASeq, we kept the fungal cultures under strict 12h:12h light-dark cycles without any ramping of light intensity. Given that ZT12 samples were collected at the transition of light-dark and that we harvested samples fairly quickly, we cannot dismiss the peaks of daily gene expression observed at the transition periods, ZT12 or ZT24, as a response to changes in light. Therefore, genes peaking before and at the light-dark transitions are most likely anticipatory peaks driven by the clock. Similar to cluster-4, we found that most genes in cluster-2 (465 genes) showed peak activity at the transition from dark to light (ZT24), whereas genes in cluster-3 (833 genes) showed anticipatory peaks between ZT20 and ZT24, a few hours before lights turned on (Fig. 1 A-B).

**Figure 2:**
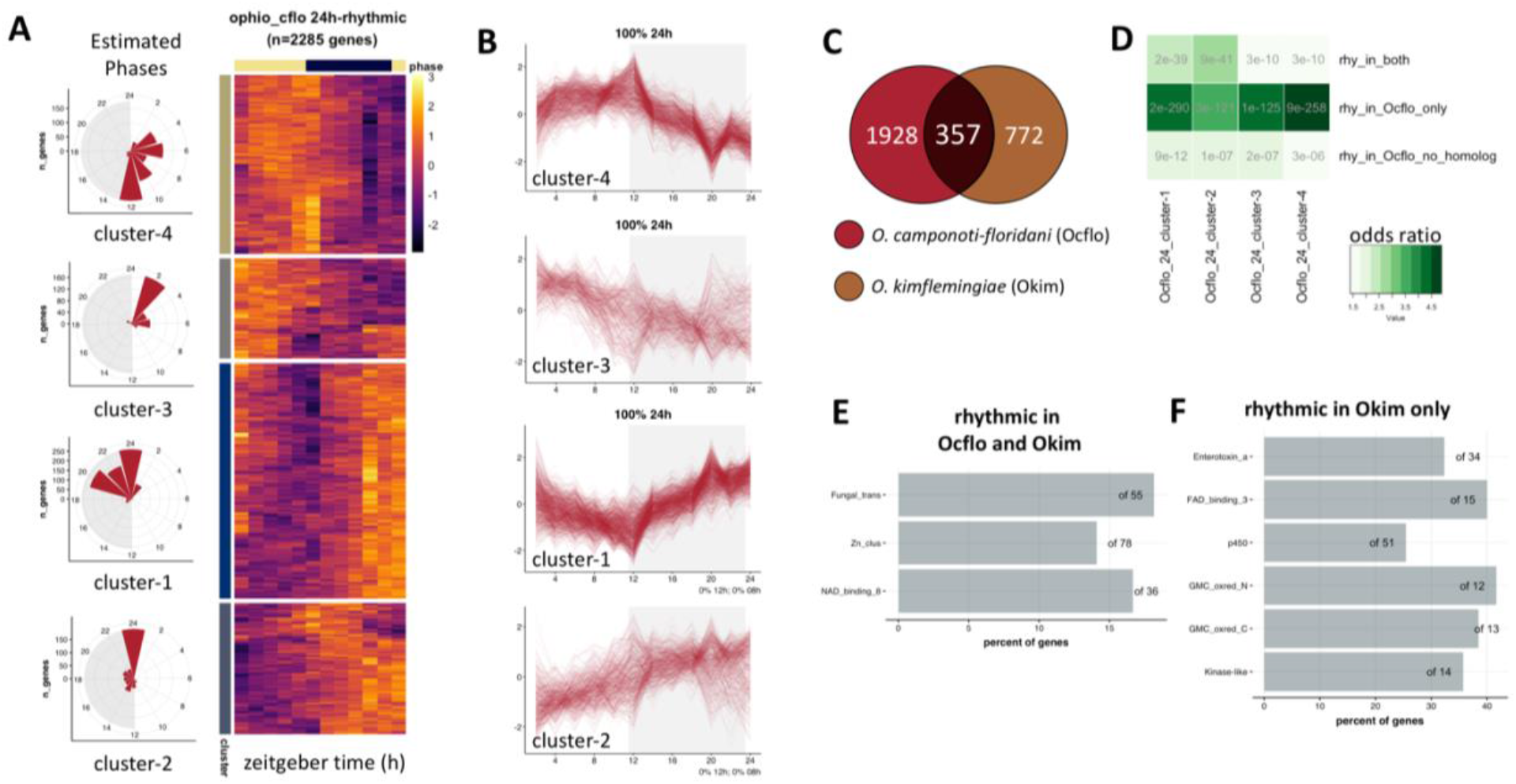
Temporal division of clock-controlled processes in Ophiocordyceps. The heatmap in panel (A) shows the daily expression (z-score) patterns of the genes identified significantly oscillating every 24h in *O. camponoti-floridani* blastospores grown in controlled liquid cultures. Each row represents a single gene, and each column represents the Zeitgeber Time (ZT) at which the sample was collected, shown in chronological order from left to right (from ZT2 to ZT24, every 2h). These 2285 rhythmic genes were hierarchically clustered into four groups based on their daily expression profiles. Each gene’s cluster identity is indicated in the cluster annotation column on the left of the heatmap. Additionally, the distribution of phases (peak time of expression) for rhythmic genes belonging to each cluster is shown on the left of the heatmap. In panel (B), we have presented the stacked zplots for all genes in a given cluster. For stacked zplots of a given cluster, each line represents the daily expression of one single gene. Panel (C) shows the overlap between the genes classified as significantly rhythmic in two *Ophiocordyceps* species (data for *O. kimflemingiae* (*Okim*) was obtained from de Bekker et al. (2021) [12] and to be consistent across the two species, rhythmicity analyses was re-run using eJTK; see Methods for more details). Panel (D) shows the results of the pairwise Fisher’s exact test that we have performed to find if certain sets of *O. camponoti-floridani* (*Ocflo*) genes significantly overlapped with either of the four distinct clusters identified in (A). The abbreviations used in the panel have the following meaning: (i) rhy_in_both = homologous genes between *Ocflo* and *Okim* classified as 24h-rhythmic in both, (ii) rhy_in_*Ocflo*_only = homologous genes classified as rhythmic in *Ocflo* but not in *Okim*, and (iii) rhy_in_*Ocflo*_no_homolog = genes that are classified as rhythmic in *Ocflo* but lack a homolog in *Okim*. Finally, panels (E) and (F), respectively, show the overrepresented pfam domains found in the set of homologous genes classified as rhythmic in both *Ocflo* and *Okim* and the homologous genes rhythmic in *Okim* but not in *Ocflo*.

Taken together, the aforementioned findings suggest that *O. camponoti-floridani* entrained its clock to the 12h:12h light-dark cycle, consistent with what was found in another *Ophiocordyceps* species [12]. Furthermore, majority of the 24h rhythmic genes of *O. camponoti-floridani* show an anticipatory peak before or at the light-dark or dark-light transitions. Light, therefore, seems to be a strong zeitgeber (entrainment cue) for the endogenous clock of *O. camponoti-floridani*. This is not surprising given that for most fungi and animals studied so far, daily fluctuations in light-dark conditions seem to be one of the prominent abiotic cues that synchronize an organism’s clock (reviewed in [37, 38]).

### Daily rhythms in gene expression – O. camponoti-floridani vs. O. kimflemingiae

Before comparing two very different fungal species with drastically different infection strategies and disease outcomes, we first compared how the clock-controlled genes and processes compared between two closely related *Ophiocordyceps* species that have both evolved with their respective ant hosts and can induce timely behavioral manipulations in their hosts. In a previous study, de Bekker et al. (2017) characterized the daily rhythms in gene expression for *O. kimflemingiae*, a closely related species of our focal *O. camponoti-floridani*, that infects and manipulates the behavior of a different carpenter ant, *C. castaneus*. We tested whether the identity and functions of clock-controlled genes in these two *Ophiocordyceps* species are similar.

Of the 8441 genes annotated in the genome of *O. kimflemingiae*, 6867 were found to have a unique homolog in *O. camponoti-floridani*. Among these homologs, 357 genes, including the fungal core clock gene *Frequency*, showed significant 24h rhythms in both *O. camponoti-floridani* and *O. kimflemingiae* (Fig. 2C). Furthermore, the overlap between rhythmic genes identified in *O. camponoti-floridani* and *O. kimflemingiae* was significant (Fisher’s exact test; odds-ratio = 1.2, p-value = 0.04). These 357 genes, seemingly conserved clock-controlled genes in *Ophiocordyceps*, were enriched for metabolic and biosynthetic processes (Fig. 2E). This finding suggests that in both *Ophiocordyceps* species, the clock drives daily rhythms in a core set of genes involved in transcriptional regulation and metabolic processes.

Next, we wanted to see if these conserved rhythmic genes showed any time-of-day preference in *O. camponoti-floridani*; are these genes primarily day-peaking or night-peaking?

To do so, we tested if the set of 357 rhythmic genes showed a significant overlap with any of the four clusters we identified for rhythmic *O. camponoti-floridani* genes (discussed in the previous section). We found that these 357 rhythmic genes (rhy_in_both) were fairly well distributed among the four clusters of *O. camponoti-floridani* rhythmic genes (Fig. 2D). Therefore, these seemingly conserved clock-controlled processes do not show a characteristic time-of-day preference, at least in *O. camponoti-floridani*.

It is known that *Ophiocordyceps*-ant interactions are highly species-specific; usually, one species of fungi can successfully manipulate the behavior of only one species of ant [7]. However, it is not known how differences in the fungal clock functioning, especially the identity and function of the clock-controlled genes, play a role in such host-specificity. To explore this possibility, we compared the homologous genes that showed significant daily rhythms in one species but not the other. Given that in the previous transcriptomics study, *O. kimflemingiae* samples were collected every 4h over a 48h period [12], as compared to every 2h over a 24h period for *O. camponoti-floridani* in our study, genes classified as rhythmic in *O. camponoti-floridani* but not in *O. kimflemingiae* might be due to a lower sampling resolution used for the latter. However, oscillating genes in *O. kimflemingiae* that were not significantly rhythmic in *O. camponoti-floridani* might be worth investigating since the sampling resolution, and sequencing depth we used for *O. camponoti-floridani* should have allowed us to identify more of the cycling genes [39].

We found 772 genes that were classified as rhythmic in *O. kimflemingiae* but not in *O. camponoti-floridani*. These genes showed significant overrepresentation in several pfam domains, including heat-labile enterotoxin, cytochrome p450, and kinase-like protein-coding genes (Fig. 2F). In summary, our findings seem to suggest that although key biological processes (transcription and metabolism) are under clock control in both *Ophiocordyceps* species, species-specific differences exist in the repertoire of clock-controlled genes and processes, some of which (e.g., enterotoxins) are hypothesized to be important for *Ophiocordyceps’* ability to successfully manipulate its specific ant host [6, 12].

### Daily rhythms in gene expression - O. camponoti-floridani vs. B. bassiana: Part One

To compare the two fungal species, we first identified the one-to-one orthologs between *O. camponoti-floridani* and *B. bassiana*. Of the 7455 protein-coding genes annotated in the *O. camponoti-floridani* genome and 10364 genes annotated in the *B. bassiana* genome, 5274 genes showed a one-to-one orthology. Of the 9006 genes expressed in *B. bassiana* grown in liquid media, 1872 genes displayed a significant 24h rhythm in daily expression, among which 701 oscillating genes lacked an ortholog in *O. camponoti-floridani*. This was similar to *O. camponoti-floridani*; 485 of the 2285 rhythmic genes did not have an ortholog in *B. bassiana*.

To characterize the differences in clock-controlled genes and processes among these two fungal species, we explored the identity and function of the non-orthologous rhythmic genes in either species. The 485 non-orthologous rhythmic genes in *O. camponoti-floridani* were enriched in heat-labile enterotoxin genes and methyltransferases (Fig. 3). In comparison, the set of rhythmic *B. bassiana* genes that lack an ortholog in *O. camponoti-floridani* showed significant enrichment for fungal-specific transcription factors: genes containing both, the catalytic domain of alcohol dehydrogenase and zinc-binding domain, beta-lactamases, NACHT domain-containing genes, and zinc finger containing transcription factors (Fig. 3).

**Figure 3.**
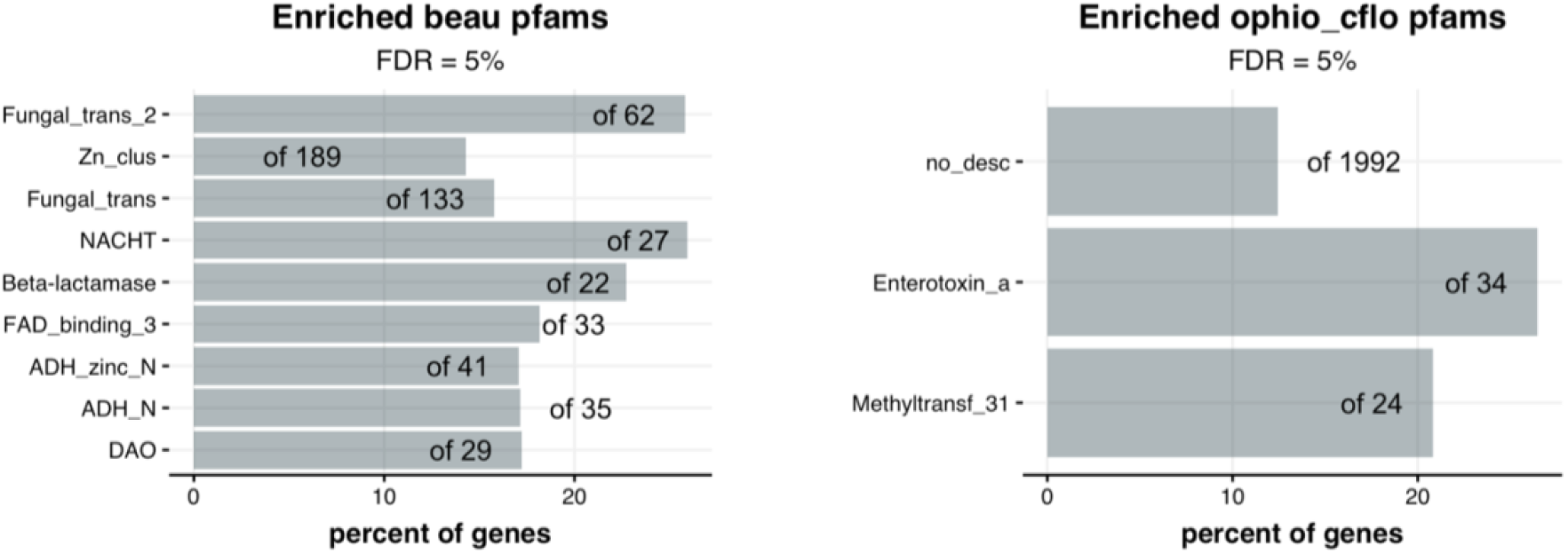
Functions of non-orthologous rhythmic genes in O. camponoti-floridani and B. bassiana. The overrepresented pfam domains found in the subset of rhythmic genes in (left) *B. bassiana* and (right) *O. camponoti-floridani* that lack a one-to-one ortholog in the other species are shown. The bars and numbers in the enrichment plot have the same meaning as before.

Beta-lactamase enzymes, usually produced by bacteria, are well known in enabling bacterial resistance against beta-lactam antibiotics such as penicillin (reviewed in [40]). Several fungal species are also known to encode beta-lactamases [40]. The functions of fungal lactamases seem more diverse than mere resistance against beta-lactam antibiotics. For example, fungal lactamases seem to enhance the fungal detoxification of xenobiotic compounds or act as a fungal pathogenicity factor recognized by the host’s immune system [40]. The daily oscillations we find in the expression of beta-lactamase encoding genes in *B. bassiana*, therefore, could either suggest a prophylactic response to daily fluctuations in circulating xenobiotic compounds or an indicator for daily fluctuations in *B. bassiana* pathogenicity to its host.

On the other hand, the finding that rhythmic genes in *O. camponoti-floridani* that lack an ortholog in *B. bassiana* comprises mostly of enterotoxin genes lines up with previous findings that suggest a role of *Ophiocordyceps* enterotoxins in manipulating host behavior [7, 12, 17]. Additionally, the rhythmic expression of *O. camponoti-floridani* enterotoxins suggests that daily fluctuations in parasite’s enterotoxin activity can potentially elicit a rhythmic detoxification response from its host.

### Daily rhythms in gene expression - O. camponoti-floridani v. B. bassiana: Part Two

In this section, we characterize the similarities and differences in the clock-controlled gene expression of *O. camponoti-floridani* versus *B. bassiana* hoping to get a glimpse into the links between fungal clock functioning and disease outcomes. Of the 5274 orthologous genes between *O. camponoti-floridani* and *B. bassiana*, 1790 genes in *O. camponoti-floridani* and 1171 genes in *B. bassiana* were 24h-rhythmic; 433 were classified as rhythmic in both, and 2519 genes were rhythmic in at least one fungal species. Since our goal was to identify synchronized changes in the daily expression of rhythmic gene clusters across the two species, we focused our further analyses on all 2519 genes that were significantly rhythmic in at least one of the two fungal species (Fig. 4A). We classified the 2519 genes into four clusters via hierarchical clustering to identify potentially day, night, dusk, and dawn peaking genes.

**Figure 4.**
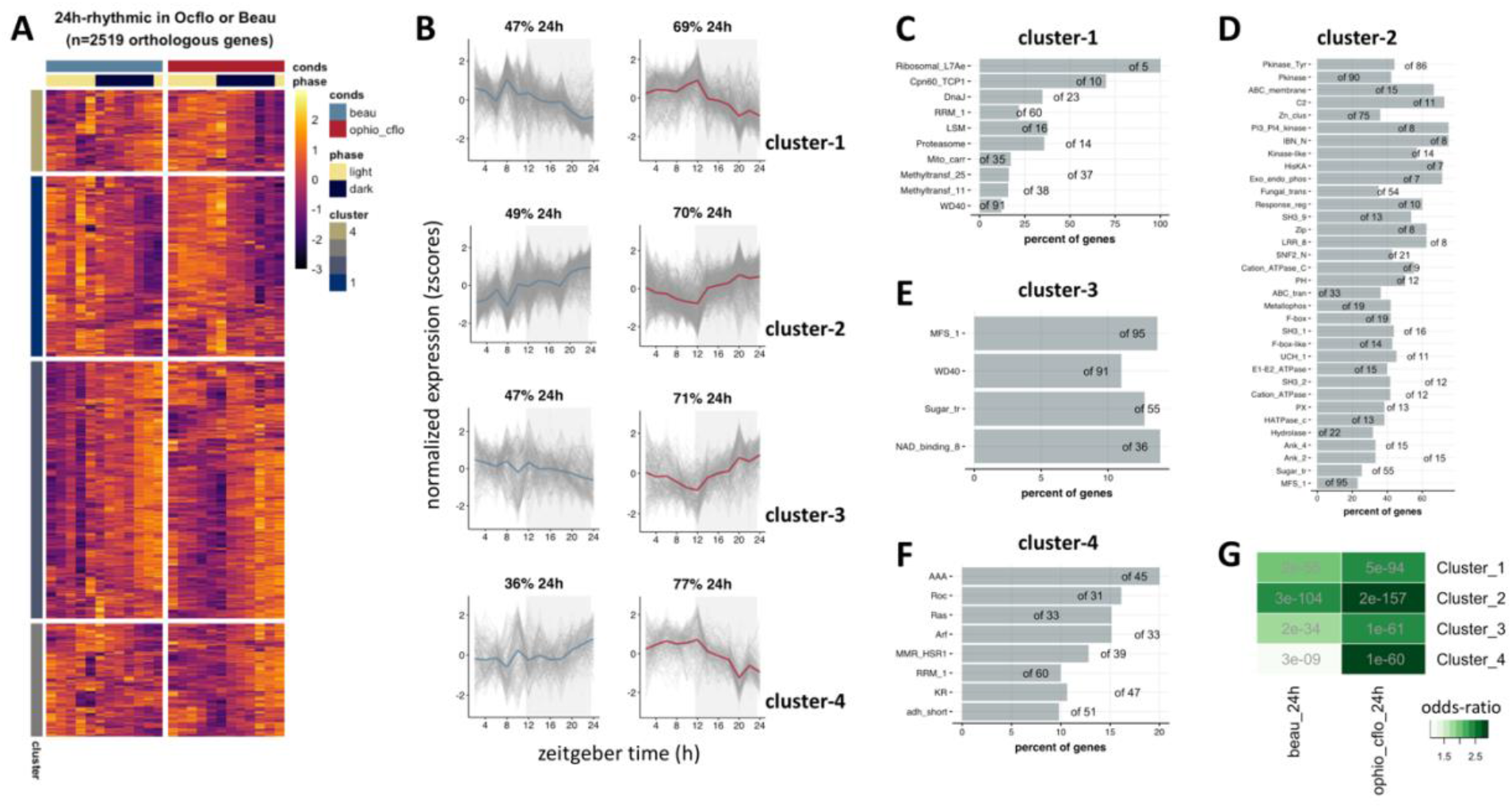
Orthologous rhythmic genes in O. camponoti-floridani and B. bassiana show species-specific differences in daily timing. The heatmap in panel (A) shows the daily expression (z-score) patterns of the orthologous genes between *O. camponoti-floridani* and *B. bassiana* that were classified as significantly 24h-rhythmic in either *O. camponoti-floridani* or *B. bassiana*. Each row represents a single gene, and each column represents the Zeitgeber Time (ZT) at which the sample was collected, shown in chronological order from left to right (from ZT2 to ZT24, every 2 h). These 2519 genes were hierarchically clustered into four groups based on their daily expression profiles in both the fungi. Each gene’s cluster identity is indicated in the cluster annotation column on the left of the heatmap. In panel (B), we have presented the stacked zplots showing the daily expression of the orthologous genes in a given cluster for *B. bassiana* and *O. camponoti-floridani*. For a given cluster, the solid-colored line represents the median daily expression of all genes in that cluster. For each stacked zplot, the number on top indicates the percentage of genes in that cluster that were classified by eJTK as significantly rhythmic as per our p-value threshold (p < 0.05). Panels (C) through (F) show the overrepresented pfam domains found in each of the four clusters identified in (A). The bars and numbers of the enrichment plot have the same meaning as before. Finally, in panel (G), we report our results of pairwise Fishers’ exact test that we performed to check which of the four clusters identified in (A) show a significant overrepresentation for genes identified as rhythmic in *B. bassiana* (beau_24h) and *O. camponoti-floridani* (ophio_cflo_24h).

Two primary observations stood out from comparing the daily gene expression of these four clusters in the two fungal species. First, the orthologous genes in cluster-1 and cluster-2 displayed a similar pattern of daily oscillations in the two fungal species, either peaking in the day-time or in the night-time. Genes in Cluster-1 showed a putative day-time peak around ZT08-Z12 in both fungi, whereas Cluster-2 peaked during the subjective night around ZT20-24. Second, genes in cluster-3 and cluster-4 showed a reversal in their phase of daily expression in the two fungal species, as is evident from Fig. 4B. Cluster-3 genes in *O. camponoti-floridani* showed a night-time peak of daily expression; the same genes in *B. bassiana* showed a day-time peak. Similarly, Cluster-4 genes showed a day-time peak in *O. camponoti-floridani*, but a night-time peak was observed for *B. bassiana*. This difference in the time-of-day of peak activity – which we call differential rhythmicity – of the same clock-controlled genes might explain, at least partly, the differences we observe in the two fungi’s life history, mode of operation inside the ant host and the eventual disease outcome. Therefore, as a next step, we explored the identity of these differentially rhythmic genes and the molecular functions they perform in fungi.

The differentially rhythmic genes in Cluster-3 showed overrepresentation for genes encoding Major Facilitator Superfamily (MFS_1 domain) of membrane proteins, some of which function as glucose transporters (Sugar_tr domain), including a probable aflatoxin efflux pump *AFLT* and an MFS drug transporter. Additionally, this cluster showed enrichment for genes containing NAD(P)-binding Rossmann-like (NAD_binding_8) domain and WD-40 repeats (WD40 domain). The latter is implicated in signal transduction and transcriptional regulation [41]. Scanning through the entire list of Cluster-3 genes revealed other genes of interest, such as two copies of *protein tyrosine phosphatase* (candidate effector of parasitic manipulation of host behavior; reviewed in [2]), a *D-3-phosphoglycerate dehydrogenase* and an *alanine--glyoxylate aminotransferase* (putative regulators of ant behavioral plasticity; [42]), and *arrestin* (involved in photoreceptor maintenance and olfactory perception in *Drosophila*).

In comparison, Cluster-4 showed enrichment for genes containing ATPase associated with diverse cellular activities (AAA domain), which included several DNA replication factors, a DNA helicase, a chromosome transmission fidelity protein, and a cell division control protein. Also overrepresented in this cluster were NAD- or NADP-dependent oxidoreductases (adh_short domain), genes encoding GTP-binding (MMR_HSR1, Roc, Arf, and Ras domains), and RNA-binding proteins (RRM_1). This cluster of differentially rhythmic genes also contained *will die slowly* (histone modification gene in flies), *Nicotinamide N-methyltransferase* (mediates genome-wide epigenetic and transcriptional change, and also implicated in xenobiotic detoxification), *Superoxide dismutase* (detoxifies reactive oxygen species), and a *glutathione S-transferase* (GST) protein-coding gene (GSTs are involved in detoxification and hypothesized to be core regulators of insect olfaction; [43]). The differences in the time of day at which clock-controlled genes in clusters 3 and 4 peak in the two fungal parasites might be key determinants of how each disease progresses in its nocturnal ant host. Future studies focused on functionally testing the role of a few of these differentially rhythmic fungal genes, especially those linked to behavioral plasticity in ants, will be important next steps.

### Differentially expressed O. camponoti-floridani genes during infection

Using time-course gene expression data generated in a separate study [9], we identified *O. camponoti-floridani* genes that showed significantly different expression levels, at the halfway mark in disease, while growing inside the ant head as compared to growth outside the host, in liquid culture. At the halfway mark in disease (referred to as “infection” from here on), we found that 318 genes were upregulated, and 395 genes were downregulated in *O. camponoti-floridani* as compared to liquid culture controls (fold-change ≥ 2, q-value < 0.05) (Fig. 5A).

**Figure 5.**
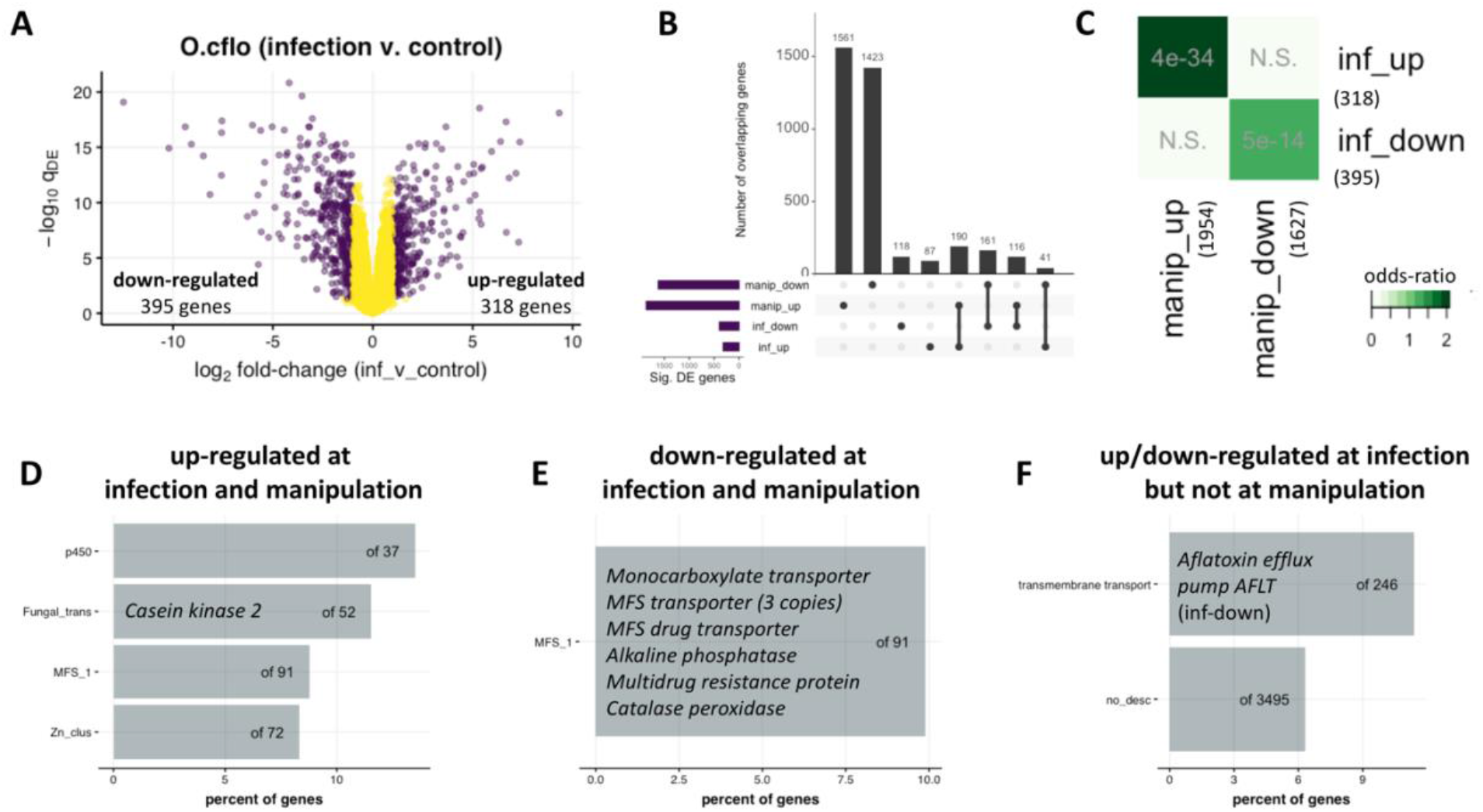
Differentially expressed genes in O. camponoti-floridani at halfway through disease progression (infection) compared to late-stage disease (manipulation). The volcano plot in panel (A) shows the results of differential gene expression analysis for *O. camponoti-floridani* sampled from within the host ant’s head halfway through disease progression (infection), compared to blastospores grown in liquid media outside the host. Panel (B) shows the intersections between four sets of *O. camponoti-floridani* DEGs (manip_down/up represents genes down/upregulated in *O. camponoti-floridani* at the manipulated biting stage as compared to controls, and inf_down/up represents genes down/upregulated in *O. camponoti-floridani* at infection as compared to controls). Panel (C) shows the results from the pairwise Fisher’s exact test we performed to check for significant overlap between the four sets of DEGs discussed in (B). In the next three panels, we show the overrepresented pfam domains we found in the set of genes (D) upregulated at both *O. camponoti-floridani* infection and manipulation, as compared to controls, (E) down-regulated at both *O. camponoti-floridani* infection and manipulation, as compared to controls, and (F) up or down-regulated at infection, but not differentially expressed at manipulation, as compared to controls. The bars and numbers of the enrichment plots have the same meaning as before. The text overlaid on top of the bars highlights some of the *O. camponoti-floridani* genes that contributed to the respective pfam enrichments.

In comparison, a previous study by our lab quantified gene expression at the final stages of the disease (referred to as “manipulation” from here on), when infected ants bit into a substrate with locked jaws before death. The previous study found that 1867 genes were upregulated, and 1625 genes were downregulated in *O. camponoti-floridani* at manipulation versus time-matched liquid culture controls (Fig. 5B; manipulation data obtained from [6]). Around 60% (190) of genes upregulated at infection were also upregulated at manipulation, whereas 41% (161) of genes downregulated during infection also were downregulated at manipulation; both the overlaps were significant (Fig. 5C).

To try and understand the role of differentially expressed genes (DEGs) in mediating the characteristic lifestyle of *O. camponoti-floridani* as it grows inside the ant host, we decided to look at three biologically relevant subsets of these DEGs, (1) genes that are significantly up- or down-regulated at infection as well as manipulation, (2) genes that are up/down-regulated only at infection but not at manipulation, and (3) genes that are downregulated at infection but upregulated at manipulation.

The genes that showed a consistent up/down-regulation at both infection and manipulation are likely involved in the general growth and survival of the fungus inside the host. For genes upregulated at both infection and manipulation, we found an overrepresentation of fungal-specific transcription factors, including the clock gene *Casein kinase 2* (Fig 19D). *Ck2* seems to play a key role in the temperature compensation of fungal [43] and plant [44] circadian clocks, which enables the cellular oscillator to function normally under different temperatures. In flies, the catalytic subunit of CK2 is required for maintaining the near 24h periodicity of the circadian clock [45]. Furthermore, the clock protein CK2 also plays an important role in mediating an organism’s response to ultraviolet light (**REFS**). It has previously been shown that several manipulating parasites, including *O. camponoti-floridani*, affect the host’s response to light, usually turning the infected hosts into light-seekers [46, 47]. Therefore, a next step might be to explore the role of CK2 as a candidate parasitic effector in modulating the host’s behavioral responses to light and temperature. In addition to transcription factors, the *O. camponoti-floridani* genes upregulated at infection and manipulation were also overrepresented for cytochrome p450 genes (oxidoreductase activity), fungal transcriptional regulatory proteins involved in zinc-dependent binding of DNA (Zn_clus domain), and membrane proteins belonging to major facilitator superfamily (transmembrane transport activity). The latter – the MFS family of membrane proteins – was also overrepresented in fungal genes down-regulated throughout the disease, both at infection and at manipulation. These downregulated MFS membrane proteins contained several transporters of interest (e.g., an *MFS drug transporter* and a *Multidrug resistance protein*). In addition, the genes downregulated throughout the disease also contained two putative *enterotoxins* and a copy of the *protein tyrosine phosphatase*.

Next, we reasoned that the second set of 205 genes that are differentially expressed (118 upregulated and 87 downregulated) only at infection but not at manipulation might be important for maintaining parasite-host homeostasis, maybe via temporal synchronization of biological functions, to ensure that infected ant host can survive the long incubation period necessary for *O. camponoti-floridani* to complete its life cycle. These 205 genes were enriched for transmembrane transport activity. They contained the *Aflatoxin efflux pump AFLT* (downregulated at infection) (Fig. 4F). This set of DEGs contained several other toxin-related genes: three enterotoxins (two *heat-labile enterotoxins* were upregulated, and a *putative enterotoxin* was downregulated), a *protoplast regeneration and killer toxin resistance protein* (downregulated), a *gliotoxin biosynthesis protein GliK* (upregulated). Gliotoxin, a known fungal virulence factor, has been implicated in assisting host tissue penetration [48]. It remains to be seen if a higher gliotoxin production at the halfway mark in the disease might be necessary for efficient delivery of the fungal enterotoxins into the host tissues, especially in the brain. Therefore, further research on the role of toxins in *O. camponoti-floridani*-induced manipulation of ant behavior would benefit from including gliotoxin as a candidate fungal effector.

This subset of around two hundred genes that were differentially expressed only at infection, but not at manipulation, contained two clock genes: *Frequency* (upregulated) and *Vivid* (synonymous to *Envoy*; downregulated). An up-regulation of the core fungal clock gene, *Frequency*, at the halfway mark in disease is intriguing. Even though *Frequency* expression show 24h rhythm in liquid culture, the temporal patterns of *Frequency* expression might or might not be rhythmic during infection. But either of the two possibilities is of interest: loss of 24h rhythms in *Frequency* expression or an increase in its amplitude of daily expression. Another important component of the fungal circadian clock, *Vivid* is essential for sensing and adapting to light in the model fungi *Neurospora crassa* [49] [50]. Light regulates various fungal processes, including phototropism, secondary metabolism, and pathogenicity (reviewed in [50]. It remains unclear to what degree the light-seeking behavior of manipulated ant hosts is an extended phenotype of the light-seeking fungal parasite growing within. But our findings suggest that it is possible.

Finally, the third set of genes that are downregulated at the halfway mark but upregulated at manipulation likely contains fungal effectors that are necessary to successfully manipulate host behavior at the final stages of the disease, e.g., death-grip, but harmful to the host otherwise. We did not find any enriched pfams or GO terms for these 43 differentially expressed genes. Although not enriched for a specific biological process, these genes might function as key regulatory components in the fungal gene expression network. In the next section, we map the several genes of interest that we have identified so far onto the gene co-expression network of *O. camponoti-floridani* and discuss the regulatory changes in the parasite’s daily gene expression that might be necessary to successfully grow inside the ant host for an extended period.

### Changes to the gene expression network of O. camponoti-floridani during infection

In this section, we took a systems approach to (1) build a daily gene co-expression network (GCN) of *O. camponoti-floridani* as observed in control (culture) conditions, (2) annotate the *O. camponoti-floridani* GCN by identifying the location of differentially rhythmic genes in the GCN, especially genes that show evidence of differential rhythmicity between *O. camponoti-floridani* and *B. bassiana* (Clusters 3 and 4; Fig. 4), and (3) identify which modules of the *O. camponoti-floridani* GCN show a drastic change in their expression (differential expression) at the halfway mark in its disease progression.

The construction and annotation of the gene co-expression network (GCN) of *O. camponoti-floridani* were done as per the protocol detailed in de Bekker and Das (2022). We built the GCN of *O. camponoti-floridani* using the 6874 genes that were expressed (≥1 FPKM) for at least half of all sampled time points in their blastospore stage grown in controlled liquid media (i.e., “controls”). We wanted to use the gene expression network of controls as the background to see what changes occur in the parasite’s network during infection. These 6874 *O. camponoti-floridani* genes were clustered into 16 modules based on their degree of co-expression and named accordingly (OC1 to OC16). The global connectivity patterns of the *Ophiocordyceps* GCN are shown in Figure 6A, where nodes represent the modules or clusters of highly co-expressed genes (Kendall’s tau-b correlation ≥ 0.7), and edges between modules indicate a similarity of module-module expression (Kendall’s tau-b correlation of at least 0.6 between a module’s eigengene expression with another).

**Figure 6.**
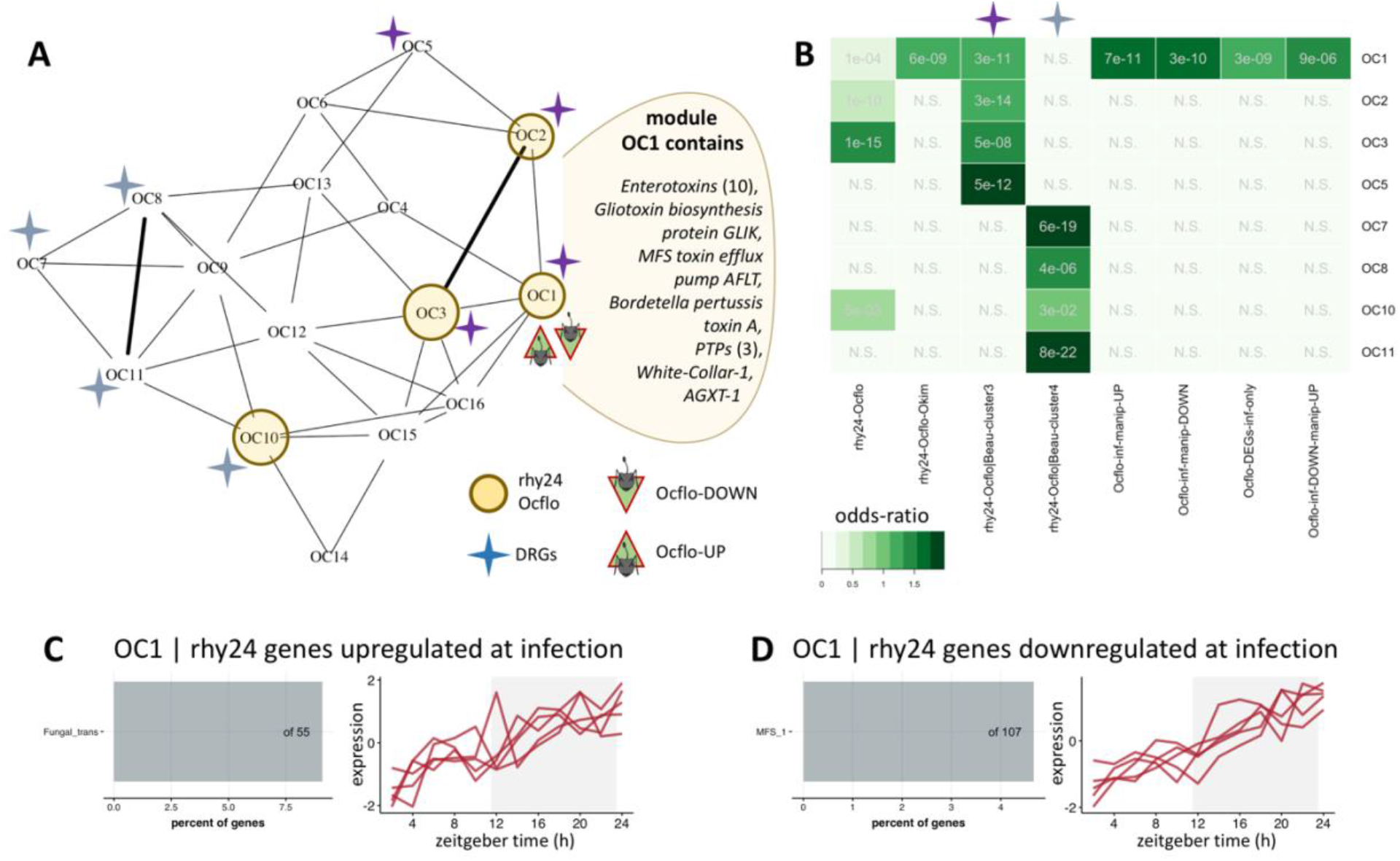
Annotated gene co-expression network of O. camponoti-floridani identifies a single cluster that potentially regulates the parasitic processes necessary to manipulate host behavior. Panel (A) shows the annotated gene co-expression network (GCN) of the manipulating fungal parasite *O. camponoti-floridani* (*Ocflo*) as observed during growth in controlled liquid media outside the host. The annotations in the network show the different modules of interest that we have identified as being putatively important for the interplay of fungal rhythmicity, disease-associated plasticity, and its ability to manipulate host behavior. The nodes represent modules of highly co-expressed genes that are clustered together, and the edges represent the co-expression of connected modules. Thick edges indicate module-module correlations ≥ 0.8, thinner edges indicate correlations between 0.6 and 0.8, and no edges indicate correlations <0.6. The abbreviations have the following meaning: (i) rhy24-*Ocflo* indicates modules that show an overrepresentation of *O. camponoti-floridani* genes that were classified as significantly rhythmic in controls, (ii) DRGs indicate the modules that show an overrepresentation of orthologous genes, between *O. camponoti-floridani* and *B. bassiana*, that show differential rhythmicity between the two species (clusters 3 and 4, Fig. 4). The two colors of the DRG symbol indicate overrepresentation for different clusters of the differentially rhythmic genes between *O. camponoti-floridani* and *B. bassiana*, as indicated in panel (B), and (iii) *Ocflo*-DOWN/UP identifies the one module in *O. camponoti-floridani* GCN that shows significant overrepresentation for several subsets of disease-associated differentially expressed genes in *O. camponoti-floridani* as compared to controls. Panel (B) shows the results of the pairwise Fisher’s exact test to identify the modules of interest in (A). Panel (C) shows the pfam domain overrepresented in the *O. camponoti-floridani* genes belonging to module OC1 that were identified as rhythmic in control conditions but upregulated at halfway through infection (referred to as “infection”). Also shown alongside is the daily expression of these *O. camponoti-floridani* genes in controls, as stacked zplots. In panel (D), we show the same information provided in (C) but for *O. camponoti-floridani* genes in module OC1 that were classified as rhythmic in controls but showed a downregulation at infection.

To understand how the genes of interest we have identified so far regulate each other, we annotated the *O. camponoti-floridani* GCN by identifying where our genes of interest are located in the network. To annotate the *O. camponoti-floridani* GCN, we identified the module(s) that showed a significant overlap with *O. camponoti-floridani* genes displaying (1) robust 24h-rhythms in *O. camponoti-floridani* controls (rhy24h-*Ocflo*), (2) genes classified as significantly rhythmic in both *O. camponoti-floridani* and *O. kimflemingiae* (rhy24-*Ocflo*-*Okim*), (3) two clusters of genes that showed differential rhythmicity between *O. camponoti-floridani* and *B. bassiana* (rhy24-*Ocflo*|Beau-cluster3 and rhy24-*Ocflo*|Beau-cluster4, Fig. 4), (4) genes that were significantly upregulated at infection as well as manipulation (*Ocflo*-inf-manip-UP), (5) genes that were significantly downregulated at infection as well as manipulation (*Ocflo*-inf-manip-DOWN), (6) genes that were differentially expressed only at infection but not at manipulation (*Ocflo*-DEGs-inf-only), and (7) genes that were downregulated at infection but upregulated at manipulation (*Ocflo*-inf-DOWN-manip-UP).

We found that most of the 24h oscillating genes in *O. camponoti-floridani* controls were located in four of the sixteen modules (OC1, OC2, OC3, and OC10; Fig. 6A-B). To validate these module annotations, we looked at the identity of the genes in these putatively rhythmic modules. As expected, we found that they contained known fungal clock components. Module OC1 contained the two core fungal clock genes, *white collar 1* and *white collar 2*, that form the positive arm of the transcription-translation feedback loop (TTFL). However, two of the other fungal core clock genes, *frequency* and *vivid*, that make up the negative arm of TTFL were located in module OC8, which did not show an overrepresentation of clock-controlled genes. Module OC8, however, showed an overrepresentation of genes identified as differentially rhythmic between *O. camponoti-floridani* and *B. bassiana* (rhy24-*Ocflo*|Beau-cluster-4; Fig. 6A-B). In addition to module OC8, the differentially rhythmic genes (DRGs) between the two fungal species were also overrepresented in six other modules, which included all four rhythmic modules (OC10, OC1, OC2, and OC3; Fig. 6A-B). Therefore, it appears that the same fungal genes that show the opposite pattern of daily expression in the two entomopathogens (e.g., day-peaking in *O. camponoti-floridani* but night-peaking in *B. bassiana*) are fairly well distributed throughout the gene co-expression network of *O. camponoti-floridani*. Furthermore, the overrepresentation of these DRGs in all of the rhythmic modules of *O. camponoti-floridani*, some of which contain putative core clock genes, suggests that differences in the life-history strategies of *O. camponoti-floridani* and *B. bassiana* are, in part, regulated by differences in how the circadian clocks temporally segregate biological processes in these two fungal species. Future studies that functionally test the role of some of these DRGs, especially the ones found in the rhythmic modules, might uncover some of the mechanistic links between a parasite’s clock and its disease progression in the host.

Next, we mapped the different subsets of *O. camponoti-floridani* genes that showed a disease-associated differential expression. The different subsets of DEGs – upregulated and downregulated genes – mapped to only one module in the *O. camponoti-floridani* GCN: module OC1. In comparison, in the host ant’s GCN, as observed in the head and the brain, disease-associated host DEGs were consistently found in a non-rhythmic module, which was connected to the rhythmic module containing most of the genes that undergo synchronized changes in their daily expression during *O. camponoti-floridani* infections [9]. Furthermore, the host’s disease-associated DEG modules contained most of the behavioral plasticity genes (caste-associated DEGs). The findings from [9] suggest that *O. camponoti-floridani* likely targets the host ant’s molecular link between behavioral plasticity and the plasticity of its clock.

Given that we found the disease-associated DEGs in *O. camponoti-floridani* to be located in the rhythmic module OC1 that also contained fungal core clock genes (*wc1* and *wc2*) show that regulatory links between disease-associated DEGs and clock-controlled processes exist in both the manipulating parasite and its ant host. However, the putative regulation seems to work differently in the manipulating parasite as compared to the host. First, the host ant’s genes up and down-regulated during *O. camponoti-floridani* infection cluster separately into distinct modules; only the down-regulated module could induce changes to the rhythmic modules. Whereas in the case of the parasite, both upregulated and downregulated *O. camponoti-floridani* genes were located in only one module, the rhythmic module OC1, which contained fungal core clock components and several clock-controlled genes.

Although it is unclear what the function of having all of these parasitic genes of interest in one rhythmic module might be, one could argue that the co-occurrence likely allows the parasite to rapidly adapt its clock-controlled processes in response to changes in its local environment, such as changes in host physiology as the disease progresses. Whether this co-occurrence has an adaptive value for the parasite’s ability to manipulate host behavior remains to be seen. However, we did find evidence suggesting that, at the very least, module OC1 plays a role even in the final stages of *O. camponoti-floridani* infection during the manipulated biting. When we mapped all the *O. camponoti-floridani* DEGs at manipulated biting (Will et al., 2020) onto our GCN, we found that the upregulated genes again were overrepresented in module OC1, while the down-regulated genes were primarily located in module OC3 and OC1. As such, it appears that during the final stages of the disease, at manipulated biting, the differentially expressed genes in *O. camponoti-floridani* show a host-like modular separation: the fungal up and down-regulated genes are no longer located in the same module. This hints at the possibility that the gene co-expression network of *O. camponoti-floridani* might undergo substantial rewiring during the final stages of the disease, as compared to the halfway mark.

Thus far, we have shown that a single cluster in the gene expression network of *O. camponoti-floridani*, module OC1, contains not only genes that are under clock control but also genes that are differentially expressed in the parasite at the halfway mark as well as in the final stages of the disease during active manipulation. We wondered, therefore, if such a regulatory gene cluster might be a hallmark of behavior-manipulating parasites, at least in the genus *Ophiocordyceps*. To answer this question, we located the genes that were classified as rhythmic in both *O. camponoti-floridani* and *O. kimflemingiae*. Again, the only module that showed overrepresentation for these conserved rhythmic genes was module OC1. Therefore, the results suggest that behavior-manipulating fungi in the genus *Ophiocordyceps* might have evolved a regulatory cluster that connects the functioning and output of the circadian clock to its ability to successfully infect an ant host.

Given that OC1 seems important to the infection biology of the hemibiotroph *Ophiocordyceps*, we further investigated the function of some putative fungal effectors that might be mediating the crosstalk between disease-associated differential expression and changes to clock-controlled rhythms. Of the 1144 genes in module OC1, 42 showed daily rhythms in *O. camponoti-floridani* controls and an upregulation during infection (the halfway mark in the disease). These 42 genes were overrepresented for fungal-specific transcription factors, including the nitrogen assimilation transcription factor *nirA*, which functions as a regulator of nitrate assimilation in fungi [51] (Fig. 6C). In the dinoflagellate *Lingulodinium polyeadra*, extracellular nitrate has been shown to act as a light-independent zeitgeber, and in turn, the clock regulates nitrate metabolism via control of nitrate reductase concentrations [52, 53]. Such clock-controlled nitrate metabolism has also been found in green algae, whereas in plants, nitrate transport seems to be under clock control (reviewed in [54]). More relevant to our findings, rhythmic nitrate metabolism has been observed in an *frq*-less *Neurospora* in which the transcription-translation feedback loop (TTFL) was dysfunctional, suggesting that rhythms in nitrate metabolism are not dependent on the fungal TTFL. Given that we observed a significant differential expression of the clock components *frq* (upregulated) and *vvd* (downregulated) during infection, it is possible that the *O. camponoti-floridani* might be employing non-TTFL means to maintain rhythmic gene expression while inside its host, at least at the halfway mark in disease. While upregulating several rhythmic transcription factors in OC1 during infection, *O. camponoti-floridani* seems to be simultaneously downregulating several rhythmic membrane transporters of the Major Facilitator Superfamily (MFS_1 domain; Fig. 6D). This functional enrichment encompassed several genes that we have already discussed previously including a *multidrug resistance protein* (GQ602_006511) and an *MFS drug transporter* (GQ602_004931). In addition to highlighting a few candidate genes in the identified module of interest, OC1, we have included complete results from our network analyses as a supplementary file (see Additional File 1 below).

In summary, comparison of the daily gene expression profile of *O. camponoti-floridani* to *B. bassiana*, a related fungal parasite that does not modify ant behavior, revealed both similarities and differences. Of the genes that were rhythmic in either of the two fungi, around half of them showed a similar temporal pattern of daily expression, either day-peaking or night-peaking, in the two fungal species. In comparison, the rest half showed different phases of daily expression in the two fungal species. This difference in the phase of the same rhythmic genes in the two fungal parasites might be contributing to the differences in their life-history strategies and disease progression in the same ant host.

Majority of the rhythmic genes in *O. camponoti-floridani* entrained to 12h:12h light-dark cycle displayed an anticipatory peak before or at the transitions between light and dark. The rhythmic genes we identified in *O. camponoti-floridani* showed significant overlap with that of a congener, *O. kimflemingiae*, that infect a different carpenter ant species, suggesting that the clock might be regulating similar biological processes across *Ophiocordyceps* species. However, we also found species-specific differences in the repertoire of clock-controlled genes, such as enterotoxin coding genes.

## CONCLUSION

This comparative study provides a first glimpse at the links between the circadian clocks of fungal entomopathogens and infectious disease outcomes for their host. We focused primarily on identifying the role of the *Ophiocordyceps* clock in modifying their host ant’s behavior. At the halfway mark in disease, when infected ants show a loss of 24h rhythms in daily foraging, several clock genes of *O. camponoti-floridani – Frequency, Vivid*, and *Ck2* – showed differential expression. Differential expression of fungal clock genes is suggestive of either a loss of 24h rhythm or an increase/decrease in the amplitude of daily expression; both possibilities suggest a change in the functioning of the *O. camponoti-floridani* clock at the halfway mark in disease, when the fungi has already grown inside the ant*’s* head but around its brain.

## SUPPLEMENTARY FILES

### Additional File 1

#### Results of the network analyses

The following CSV file contains the results from the network analysis. It includes the list of all ant genes used to build the gene co-expression network, their module identity, functional annotations, and all relevant information necessary to follow the main text.

*Link to file*:

https://github.com/biplabendu/Das_et_al_2022a/blob/master/manuscript/99_dissertation/07_Ocflo_GCN_results.csv

## Notes

### Competing Interest Statement

The authors have declared no competing interest.

### Summary of Updates

Author ORCIDs have been added. No changes have been made to the manuscript.

